# A domain-agnostic MR reconstruction framework using a randomly weighted neural network

**DOI:** 10.1101/2023.03.22.533764

**Authors:** Arghya Pal, Lipeng Ning, Yogesh Rathi

## Abstract

**Purpose:** To design a randomly-weighted neural network that performs domain-agnostic MR image reconstruction from undersampled k-space data without the need for ground truth or extensive in-vivo training datasets. The network performance must be similar to the current state-of-the-art algorithms that require large training datasets.

**Methods:** We propose a Weight Agnostic randomly weighted Network method for MRI reconstruction (termed WAN-MRI) which does not require updating the weights of the neural network but rather chooses the most appropriate connections of the network to reconstruct the data from undersampled k-space measurements. The network architecture has three components, i.e. (1) Dimensionality Reduction Layers comprising of 3d convolutions, ReLu, and batch norm; (2) Reshaping Layer is Fully Connected layer; and (3) Upsampling Layers that resembles the ConvDecoder architecture. The proposed methodology is validated on fastMRI knee and brain datasets.

**Results:** The proposed method provides a significant boost in performance for structural similarity index measure (SSIM) and root mean squared error (RMSE) scores on fastMRI knee and brain datasets at an undersampling factor of R=4 and R=8 while trained on fractal and natural images, and fine-tuned with only 20 samples from the fastMRI training k-space dataset. Qualitatively, we see that classical methods such as GRAPPA and SENSE fail to capture the subtle details that are clinically relevant. We either outperform or show comparable performance with several existing deep learning techniques (that require extensive training) like GrappaNET, VariationNET, J-MoDL, and RAKI.

**Conclusion:** The proposed algorithm (WAN-MRI) is agnostic to reconstructing images of different body organs or MRI modalities and provides excellent scores in terms of SSIM, PSNR, and RMSE metrics and generalizes better to out-of-distribution examples. The methodology does not require ground truth data and can be trained using very few undersampled multi-coil k-space training samples.

## 1 INTRODUCTION

Magnetic Resonance Imaging (MRI) has become a *sine qua non* in the fields of radiology, medicine, and psychiatry. This powerful, non-invasive imaging technology provides detailed images of the soft tissue without harmful radiation as in other modalities like CT. By design, MRI scanners provide us the patient’s anatomy in the frequency domain commonly referred to as “k-space”. The acquired k-space rows/columns are directly proportional to the quality (and spatial resolution) of the reconstructed MR image [1]. A long scan time is generally required to get good quality MR images. However, long scan times lead to motion artifacts (due to subject movement) and is an inconvenience to the patients with medical conditions such as claustrophobia, obesity, heart disease, etc. One might choose to skip acquiring a few k-space lines in order to shorten the acquisition time but this leads to the well-known “aliasing” problem as it violates the Nyquist sampling criteria [2]. In this paper, we address this problem of accurately reconstructing the MR image from undersampled k-space data by learning a nonlinear mapping from the undersampled k-space data to the MR image using a neural network model. Our methodology offers a novel perspective that provides fast multi-coil MRI reconstructed images using an untrained weight agnostic randomly weighted network (WAN) by doing a neural architectural search using the popular “strict lottery ticket hypothesis” methodology [3]. The proposed methodology can find a weight agnostic optimum network using *only* the natural and fractal images without the need for ground truth MRI data, which is a significant advantage over most deep learning state-of-the-art methods.

### 1.1 Classical methods

A classical k-space to MRI reconstruction method includes GRAPPA [4] or its variants that interpolate the missing k-space lines. These methods use the the auto-calibration or ACS lines (i.e. k-space lines from the center of the k-space) to learn kernels that can be used to interpolate the missing k-space lines. This methodology is used algorithms such as SMASH [5], VDAUTOSMASH [6], and GRAPPA and its variations [7, 8, 9]. The k-t GRAPPA method [10] takes advantage of the correlations in the k-t space and interpolates the missing data. On the other hand, sparsity promoting low rank models are based on the assumption that k-space should follow a structure that has low rank due to the limited support of the images in the MR space. The low rank assumption has been shown to be quite successful in reconstructing images as demonstrated in several existing techniques [11,12,13,14,15,16,17]. Another class of works investigate the use of compressed sensing (CS) in MR reconstruction after its huge success in the field of signal processing [18, 19, 20]. Compressed sensing requires incoherent sampling and sparsity in the transform domain (Fourier, Wavelet, Ridgelet or any other basis) for nonlinear image reconstruction. On the other hand dictionary learning based approaches such as [21, 22, 23, 24, 25] show various ways to estimate the image and the dictionary from limited measurements.

### 1.2 Deep learning methods

MR image reconstruction using deep learning, in its simplest form, amounts to learning a map G from the undersampled k-space measurement 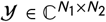, or 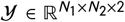 to an unaliased MR image 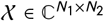, or *χ* ∈ ℝ^*N*_1_×*N*_2_×2^, where *N*_1_, *N*_2_ are the height and width of the complex valued image. In several real-world cases, higher dimensions such as time and volume are obtained and accordingly the superscripts of 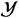 and **χ** change to 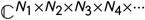. For the sake of simplicity, we will use assume 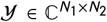 and 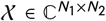. A DL method learns a non-linear function, 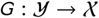 or, we can say **x**_*i*_, = *G_θ_* (**y**_*i*_) parameterized by *θ*, from a set of all possible mapping functions 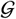. The accuracy of the mapping function can be measured using some notion of a loss function 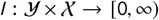. The empirical risk [26], 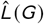, can be estimated as 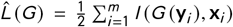 and the generalization error of a mapping function *G*(·) can be measured using some notion of accuracy measurement. Based on how we optimize the parameter *θ* in the equation **x**_*i*_, = *G_θ_*(**y**_*i*_) we can classify the DL methods into two categories: (i) the training phase methods, and (ii) the no-training phase methods.

Given the k-space measurements, the *training phase based methods* have a dedicated training phase during which the weights of a neural network are optimized in response to the estimated error given the networks prediction of MRI image and the ground truth MRI image. One such training phase based method is the AUTOMAP algorithm [27] that learns a reconstruction mapping using a network having three fully connected layers (3 FCs) and two convolutional layers (2 Convs) with an input dimension of 128 **×** 128. Similarly, several methods such as [28, 29, 30, 31] optimize the weights of a Convolutional Neural Network (CNN) for MR image reconstruction. Recently developed methods such as [32, 33, 34] optimize a Generative Adversarial Network (GAN) for MRI reconstruction. Different from these, a local-global recurrent neural network [35] uses a bidirectional LSTM that replaces the dense network structure of AUTOMAP [27] for removing aliasing artifacts in the reconstructed image. The Convolutional Recurrent Neural Networks or CRNN [36], VariationNET [37], Deep Variational Network of [38] method propose variable splitting and alternate minimisation method for MRI reconstruction.

The *no-training phase methods* solve the inverse problem using Deep Learning models that do not necessarily require a training phase to learn the optimum weights. These methods update the weights directly during test-time or in this case while performing MR reconstruction. For example, Deep Image Prior (DIP) and its variants [39,40,41] have shown outstanding performance in several computer vision tasks and have been successfully applied to perform MRI reconstruction. Similarly, the Deep Decoder algorithm [42] shows that an under-parameterized decoder network *D_θ_d__*(·) will not learn the high frequency components such as noise and can nicely approximate the denoised version of an image. The Deep Decoder network uses pixel-wise linear combinations of channels and shared weights in spatial dimensions that collectively help it to learn relationships and characteristics of nearby pixels. It has been recently understood that such advancements can directly be applied to MR image reconstruction [43]. The “Scan-Specific Artifact Reduction in k-space” or SPARK and related methods [44,45,46] show MRI reconstruction without explicit requirement of a training phase. Similar to the spirit of GRAPPA method, the robust artificial neural network for k-space interpolation (RAKI) [47] trains a CNN by using the ACS lines. Followup works such as the residual RAKI (rRAKI) [48], LORAKI [49], and sRAKI-RNN [50] methods proposed a unified framework that performs regularization through calibration and data consistency using the k-space measurements alone. The works in [45, 46] provide theoretical guarantees on recovery of the image from the k-space measurements.

The training phase methods require a long time to optimize the weights using a large dataset at training time which might leverage training time cost, dataset curation cost, human bias at the time of data collection, privacy concerns of the collected training data, etc. It is also observed that the training phase methods are not task-agnostic and perform poorly for out-of-distribution test cases such as reconstruction of brain images from a network trained on the knee dataset [51]. On the other hand, the non-training phase based methods do not require explicit training datasets but perform optimization during inference time which makes them extremely slow, requiring several minutes to process a single slice of k-space data.

In this work, we ask the question *“what is needed to design a good DL based MRI reconstruction method that addresses most of the concerns discussed above?”* To answer this question, we adopt the following two guiding principles:

- **Principal 1**: can a network structure by itself encode informative cues to solve the MR reconstruction problem from highly under-sampled k-space measurements, without changing the weight of the neural network?
- **Principal 2**: can we do away with in-vivo training datasets entirely and possibly learn the k-space to MRI image mapping from natural and synthetically generated fractal images?

To this end, we propose an untrained weight agnostic randomly weighted network (WAN) that is *different from both* the training phase based methods and the non-training phase based methods. The WAN methodology selects an optimal subnetwork from a randomly weighted dense network to perform MR reconstruction without updating the weights neither at training time nor at inference time. Our methodology does not require ground truth MRI data and shows excellent performance across domains in T1-weighted (head, knee) images from highly under-sampled multi-coil k-space measurements. The proposed methodology offers a novel perspective to learn the mapping from undersampled k-space to the MR image. We show that the proposed method outperforms the state-of-the-art (SOTA) methods such as ConvDecoder, DIP, VarNet, etc., on the fastMRI knee and brain datasets.

## 2 METHODOLOGY

### 2.1 Multicoil Accelerated MRI Reconstruction

Modern MRI scanners supports parallel acquisition that is comprised of an array of *n* overlapping receiver coils modulated by their sensitivities *S_i_*. The MRI scanner provides an MR image 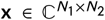 from a fully sampled k-space measurement 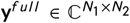 using the Fourier transform 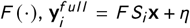, where *i* = {1,2, ⋯, *n*}. To reduce the scanning time as well as geometric distortions, the k-space is typically undersampled to acquire **y**_*i*_, coming from the *i^th^* receiver coil after applying a binary sampling mask 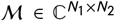 that selects a subset of k-space lines from **y**^*full*^ in the phase encoding direction. The measurement is corrupted with noise 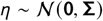 and is assumed to have a Gaussian distribution [52].

The k-space to MRI image reconstruction (for both training phase methods and no-training phase methods) broadly try to optimize the following p-norm loss function:

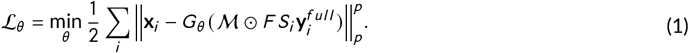

This data fidelity loss function is typically used to ensure that the estimated data is as close to the measurement as possible [37, 53]. Typically, *G_θ_* **(·)** is a neural network consisting of *m* layers *I*_1_, ⋯, *I_m_*. There are a few methods [54, 55, 56] that use m separate neural networks *G_θ,i_* (·) for each coil separately and then combine them at a later stage. While, on the other hand, there are few other methods that use a single *G_θ_* (·) network to reconstruct MRI images given all the k-space images from coils. In some cases the methodology provides the filled-up k-space given the undersampled data and we can reconstruct the image after applying an inverse fourier transform, i.e. 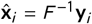. The coil-combined image can be obtained using the root-sum-of-squares (RSS) algorithm, i.e. 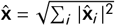.

### 2.2 Weight Agnostic Randomly Weighted Network (WAN)

In this section we asked ourselves the question; “Can a random weighted deep network structure encode informative cues to solve the MR reconstruction problem from highly under-sampled k-space measurements?” To this end, we consider the formulation in Eqn. 1, i.e. 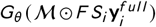, but instead of training the weight, *θ*, we performed a neural architecture search (NAS) to get a subnetwork from a randomly initialized network 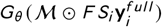. In this case, the network *G* : ℝ^*n*×*w*×*h*^ → ℝ^*n*×*w*×*h*^ has *w, h* as its width and height and *n* is the number of coils. So, our proposed methodology selects an optimal subnetwork from a randomly weighted dense network to perform MR reconstruction without updating the weights - neither at training time nor at inference time. The weight agnostic random weight is a special case of the “Lottery Ticket Hypothesis” [57] that we would refer as “Strict Lottery Ticket Hypothesis” first proposed in [3]. The lottery ticket hypothesis says that there exists a subnetwork (known as “winning ticket”) of a dense, randomly initialized, bigger neural network that when trained in isolation would reach or overcome the test accuracy of the original bigger neural network. On the contrary, the strict lottery ticket hypothesis says there exists a subnetwork of a randomly initialized network which can outperform the original network even without updating the weights of the subnetwork. Driven by strict lottery ticket hypothesis, our methodology finds a subnetwork, *G_ϕ_* (·) (the subnet is parameterized by *ϕ* and *ϕ* ⊂ *θ*) from the network, *G_θ_* (·), by dropping few weights and connections from *G_θ_* (·) using a modified version of the “edge-popup” algorithm [3] also referred to as the “edge-discarding algorithm”.

#### The Edge-discarding Algorithm

For any weight of either the fully connected or convolutional layer we associate a score value *α* ∈ *A* that decides whether or not that weight is to be included in the subnetwork. The methodology drops all the weights with low score values to find a subnetwork in the parameter space *θ* of the network *G_θ_* (·). The score update rule is given by:

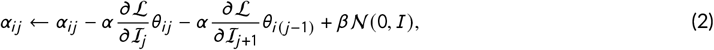

which updates the score values of the weights. Here, *θ_ij_* is output of node *i* at layer l, and 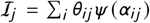 is the summed up input to node *j* at layer l + 1 from all nodes at layer l, the Gaussian noise 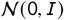 acts as a regularizer to the update rule, while *α, β* are the learning rates, and *ψ* (·) = 1 if it is above a certain threshold and 0 otherwise. The weights undergo a permutation after running for few epochs. This acts as an extra regularization to the edge-popup algorithm.

#### Learning Supermask Using Edge-discarding Algorithm

Without loss of any generality, we could say that we could use the edge-discarding algorithm to identify the plausible “supermask”, ***κ*** that acts on the weight *θ*, i.e. *ϕ* = ***κ*** ⊙ *θ*, of the network *G_θ_* (·) to get a subnetwork *G_ψ_*=_(*κ*⊙*θ*)_ (·) that would provide the best accuracy for the following loss function:

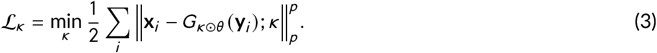

We note that getting ground truth MRI images **x** might be challenging and as such we need to modify our methodology to not require the existence of ground truth data. Consequently, we propose to learn a supermask by enforcing k-space measurement data consistency [54] in Eqn. 3 without requiring full-sampled MRI ground truth data **x**. By simply enforcing data consistency on the acquired data, we can find an optimal supermask by minimizing the following loss function:

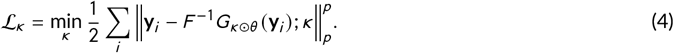

To leverage additional regularization while learning the “supermask” ***κ***, we add a *self-supervision loss* similar to [58] in Eqn. 4. To elaborate, let’s assume that we have a k-space measurement **y**. We could artificially drop a subset of k-space lines from the already undersampled data **y** and get 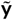. Thus, **y** is the true undersampled data obtained from the scanner while 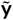 is the data obtained by further undersampling **y**. We use this 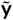 in our learning task and to enforce data consistency (serving as a pseudo ground truth). While learning the “supermask” ***κ*** we use 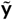> as input and perform the following optimization:

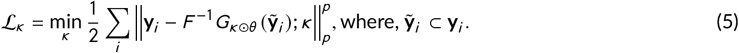

This WAN-MRI setup is motivated by the work of [**?**]. In their work however, they do not consider the random weight, edge-discarding algorithm.

### 2.3 WAN Optimization using Fractal Images

We use iterated function systems [59] (IFS) to create fractal images for training. The brain has many self-similar or fractal structures (cortical folds) and hence we hypothesize that training on synthetically generated fractal images will enable better performance of the reconstruction task at hand compared to purely natural images. To create a fractal image **x**_*f*_, we consider the sequence 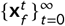 that changes the intensity of pixels by drawing dots on a black background using the random iteration algorithm proposed in [59]:

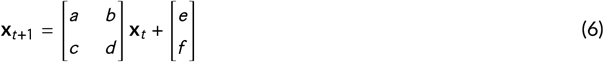

By choosing different values of [*a, b, c, d, e, f*], one could get the different shapes of fractal images that we show in Fig. 1.

**FIGURE 1.**
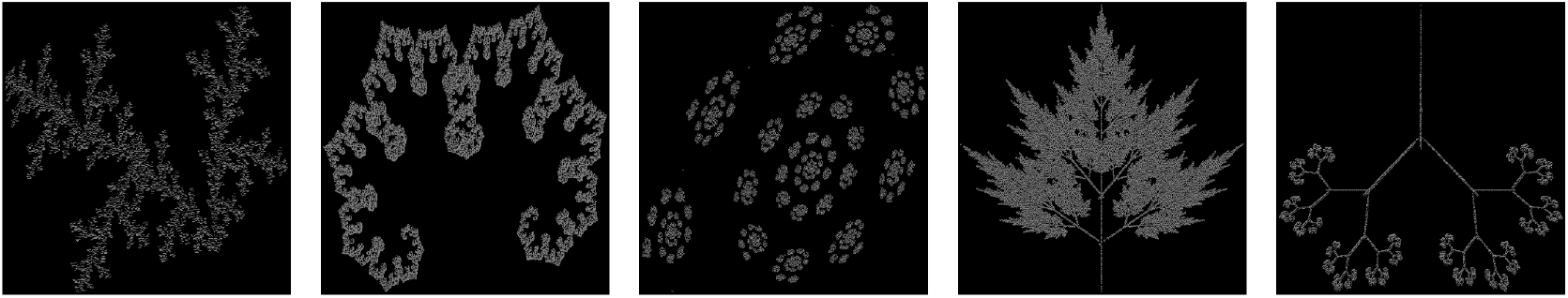
Different classes of fractral images: We demonstrate few fractral images

**FIGURE 2.**
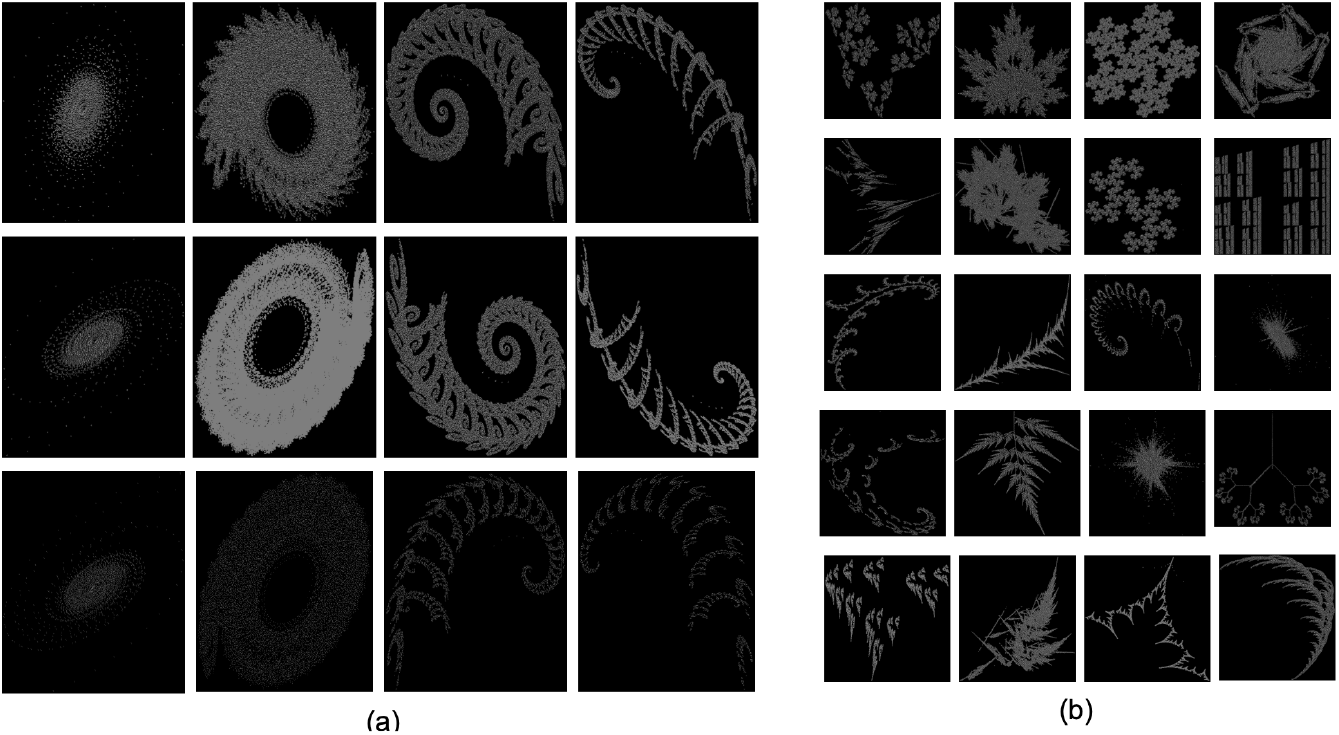
More Fractal Images. (a) We can get different images of a fractal group. We changed the orientation in second row images and the point density in third row images. (b) Different styles of fractal images can be leveraged by changing the hyperparameters of Eqn. 6.

Such fractals images, i.e. **x**_*f,i*_ could be used in Eqn. 7 to learn the supermask ***κ***:

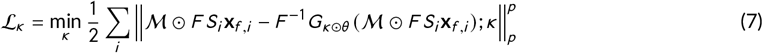

Note that, k-space of **x**_*f*_ is generated and used during training with the desired undersampling factor. In this way, our methodology leverages fractal image k-space as input and learns the supermask ***κ***. In this work, we have chosen acceleration factor *R* = 2 to get the undersampled k-space data. We have discuss more about this in Sec. 3.

### 2.4 WAN-MRI Algorithm

The overall algorithm can be summarized as follows:

- Step 1: The supermask ***κ*** is optimized using 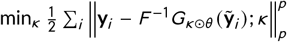, where, 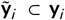, keeping random weights untouched. Here, 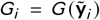 is the *i*-th output channel of the WAN. Each *G_i_* provides inferred fully sampled k-space 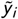.
- Step 2: We get *x_i_* = **F**^-1^ (*y_i_*) and the reconstructed MR image using the root-least-squares algorithm, i.e. 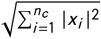.
- Step 3: Similar to ConvDecoder [54], we also enforce data consistency of the resulting image.
- Step 4: Repeat the procedure with different initializations and average the results.

## 3 EXPERIMENTS

We used the fastMRI knee and brain datasets in this work for training, testing and validation. To study the efficacy of our proposed method we compare our method with seven state-of-the-art methods such as GRAPPA, SENSE, RAKI, JMODL, VarNet, ConvDecoder, and Deep Image Prior (DIP). We used the original implementations ^1^ of GRAPPA, SENSE, RAKI, JMODL, VarNet, ConvDecoder, and Deep Image Prior (DIP). Where applicable, we always use the central 30 k-space lines to compute the training target. We treat the real and imaginary parts of the data as two distinct channels. Where applicable, the models were trained with a linear combination of *L*_1_ and SSIM loss, i.e.

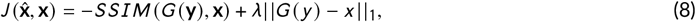

where *λ* is a hyperparameter, *G*(**y**) is the model prediction, and **x** is the ground truth. The fastMRI knee dataset consists of raw k-space data from 1594 scans acquired on four different MRI machines. We used the official training, validation and test data split in our experiments. We did not use images with a width greater than 372 and we note that such data is only 7% of the training data split. Both the 4x and 8x acceleration factors were evaluated.

We obtained the dMRI k-space measurements on a 3 Tesla (3T) Prisma MR scanner. The acquired k-space dimension of dMRI scans were (75 × 100) that we converted to dimension (100) using Projection onto Convex Sets (POCS) formalism (POCS) (corresponding to spatial resolution of 2 mm isotropic voxel size). The acquired data had 32 coils, 64 slices, and 64 gradient directions (including b=0 images). We show qualitative and quantitative comparisons of the state-of-the-art (SOTA) methods and our WAN-MRI model in Secs. 3.2 and 3.3.

Several variations of the fractal images were created by varying the hyperparameter values in Eqn. 6. In Fig 1 (a) we could generate swirl like patters by varying the values of [*a, c, e, f*] to [(0.41 to 0.81), (0.47to 0.57), (−0.86), (−0.07)]. Similarly we could get different fractal patterns shown in Fig. 1 (b) by changing the values of [*a, b, c, d, e, f*] from the numerical range [-0.01to 1.05]. All these images served as the training data for our WAN-MRI network.

### 3.1 Network Structure

We used the network architecture shown in Fig 3. The architecture has three components, i.e. (1) Dimensionality Reduction Layers comprised of 3d convolution, ReLu; (2) Reshaping Layer is Fully Connected layer; and (3) Upsampling Layers that resembles the ConvDecoder [54] network. The 3d convolution layers are comprised of (3,3,3) kernels to use (*coils, height, width*) as inputs. There are eight such 3d convolution layers that progressively reduces the input dimension and provides input to the reshaping layer. Reshaping layers are comprised of three 100 × 100 layers followed by the upsampling layers. We neither use any sort of skip connections nor batchnorms in our architecture. We tested both the stochastic gradient descent (SGD) and ADAM optimization techniques in our experiments and found that the ADAM optimization performed better than SGD. We chose a learning rate of 0.01, 1*e* - 4 as the weight decay, 0.9 for momentum and a batch size of 64. However, there is no visible change in results if we use any other forms of weight initialization. The

**FIGURE 3.**
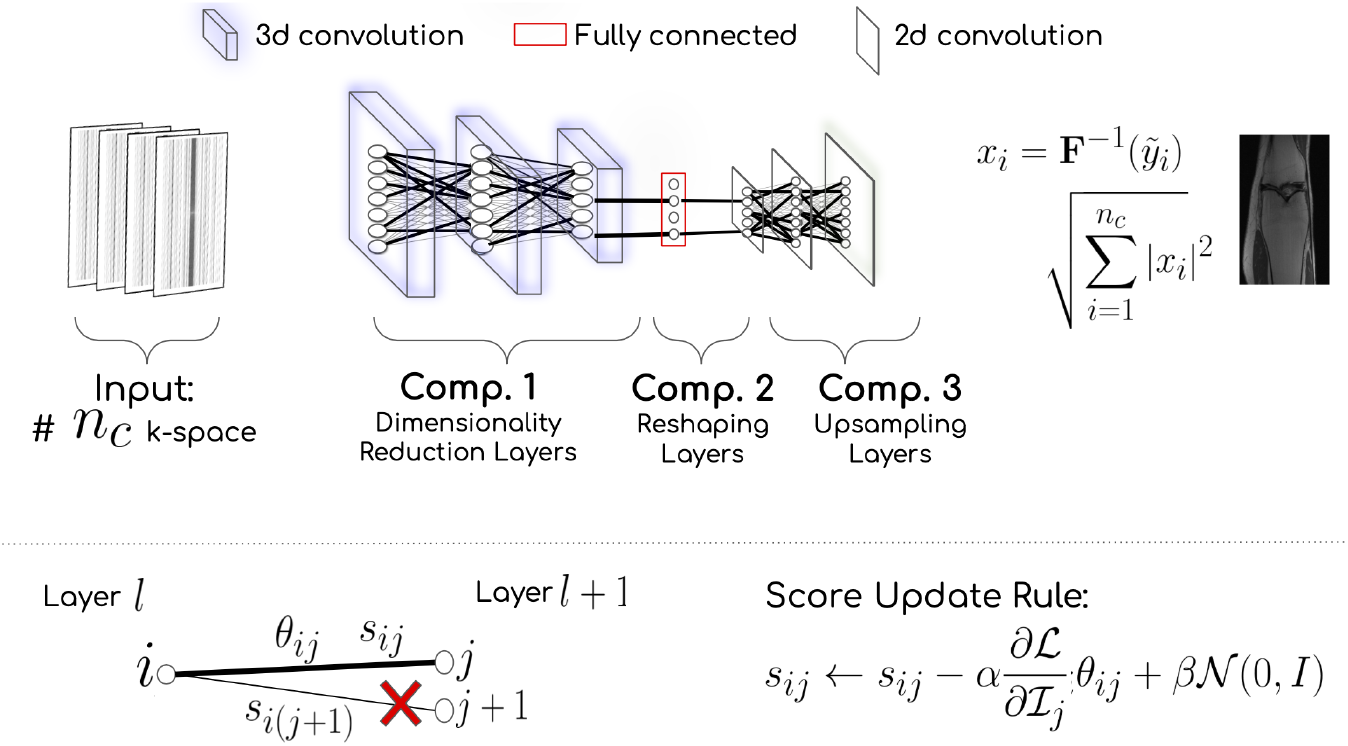
WAN architecture has 3 components, i.e. (1) Dimensionality Reduction Layers comprised of 3d convolution, ReLu, and batch norm; (2) Reshaping Layer is Fully Connected layer; and (3) Upsampling Layers that resembles the ConvDecoder [54]

### 3.2 Qualitative Results

Among the methods that we use for comparison, GRAPPA requires use of ACS lines to learn the kernel (and hence requires extra acquisition), whereas SENSE, Deep Decoder and DIP do not require any training data or ground truth data. On the other hand, RAKI, JMODL and MODL require extensive training and ground truth datasets, which potentially limits their applicability. On the other hand, the proposed WAN-MRI too does not require in-vivo MRI ground truth dataset for training.

We perform extensive evaluation on several datasets. Figs. 4 and 5 show results for R =4 undersampling on the fastMRI brain and knee datasets in respectively. The GRAPPA kernels are learned from 30 ACS lines. The learned kernels are then applied to other measurements. We note that SENSE and GRAPPA smooth a majority of the minute high frequency (edges) details that are clinically and anatomically important as seen in Figures 5 and 4. The neural network based RAKI method performs better than classical methods as seen in the values of RMSE and SSIM in Tables 1 and 2 and as is evident in Figs 5 and 4. The difference images in Figs 5 and 4 show that RAKI does less smoothing of the data. Outstanding performance of Deep Decoder and DIP advocates the importance of letting untrained neural network figure out how to perform k-space to MR image reconstructions. The JMODL method performs exceptionally well but makes heavy use of training data and the joint optimization of k-space lines and the reconstruction of MR images to get good results both for *R* = 4 and *R* = 8 as shown in Tables 2 and 1. On the other hand, the Deep Decoder and DIP methods achieve good performance using untrained networks as discussed in Sec. 1, which is advantageous as it generalizes to any MR reconstruction scenario.

**FIGURE 4.**
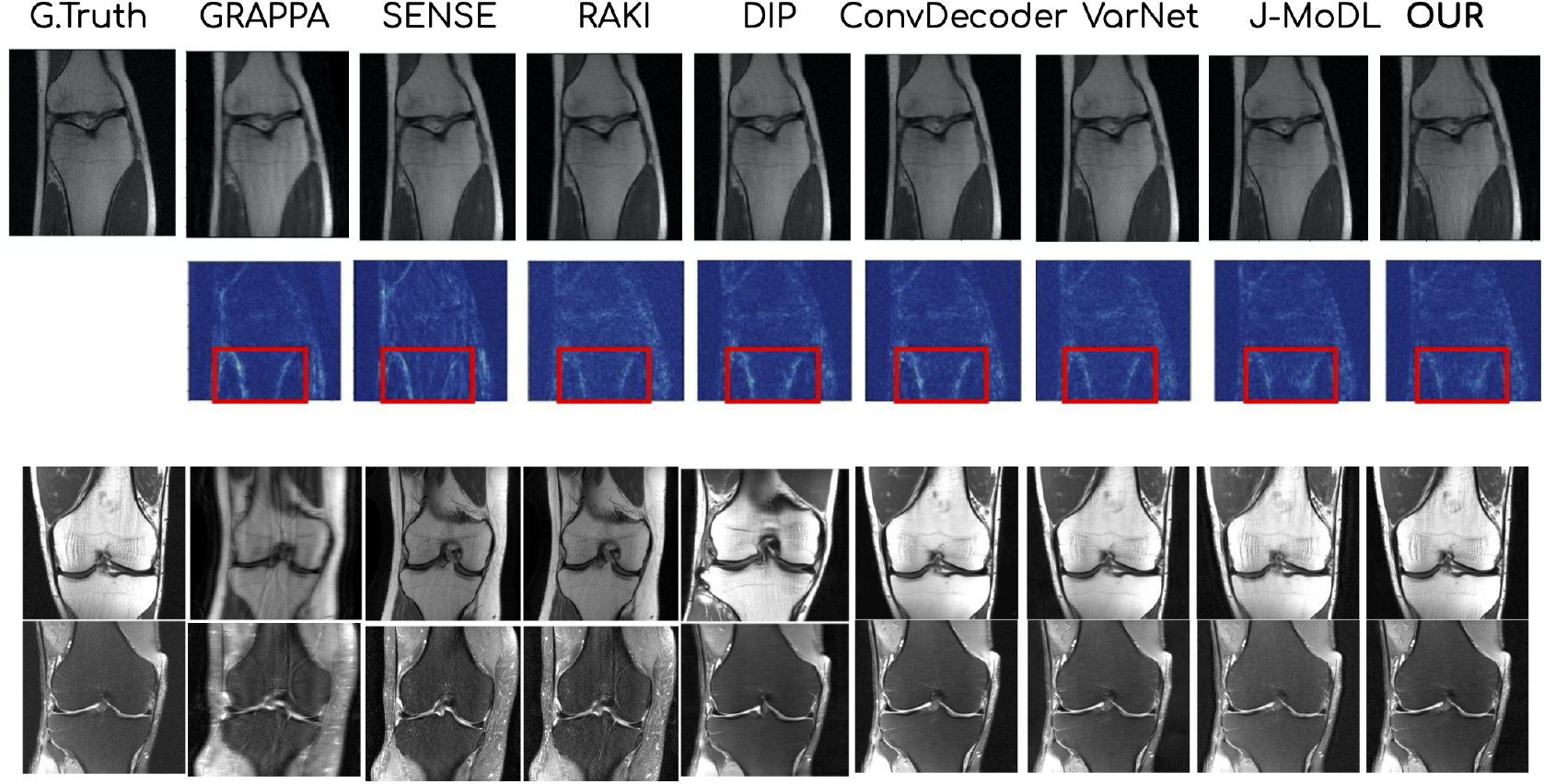
Quantitative Comparison fastMRI Knee Dataset: Quantitative comparison of reconstructions of different methods. The middle row is the difference map with ground truth and the red boxes indicate the reconstruction quality of a region. All networks were trained on the FastMRI knee training dataset.

**FIGURE 5.**
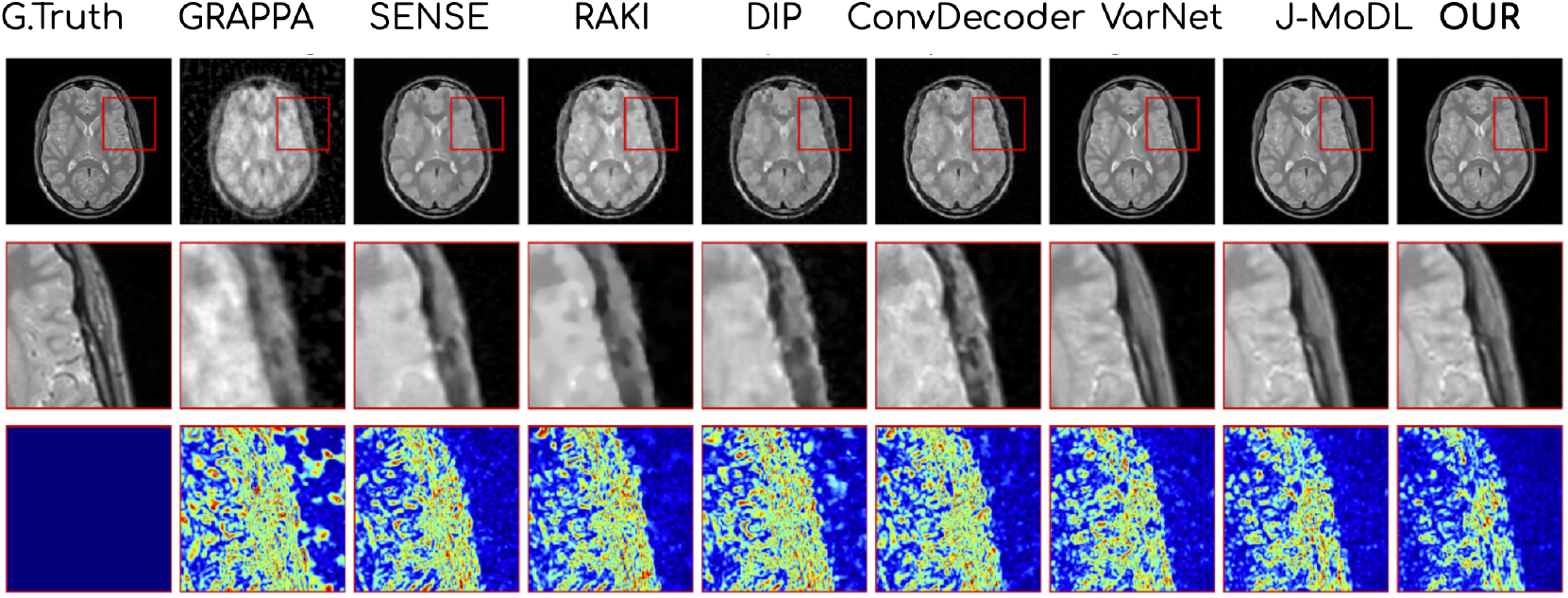
Quantitative Comparison fastMRI Brain Dataset: Quantitative comparison of reconstructions of different SOTA methods. The middle row shows a zoomed-in region. The difference map shows that our method is on par with the SOTA methods. All networks were trained on the FastMRI brain training dataset.

**TABLE 1.** fastMRI Knee: Our method outperformed all SOTA methods when our method is trained on procedural images, i.e. fractral images, natural images from ImageNet, and a small set, i.e. only 20 samples from MRI database.

|  | R = 4 |  | R = 8 |  |
| --- | --- | --- | --- | --- |
|  | RMSE | SSIM | RMSE | SSIM |
| GRAPPA | 0.0154 +/- 0.002 | 0.791 +/- 0.002 | 0.0351 +/- 0.002 | 0.642 +/- 0.002 |
| SENSE | 0.0198 +/- 0.002 | 0.806 +/- 0.001 | 0.0261 +/- 0.002 | 0.682 +/- 0.002 |
| RAKI | 0.0138 +/- 0.006 | 0.822 +/- 0.002 | 0.0221 +/- 0.002 | 0.792 +/- 0.004 |
| JMODL | 0.0055 +/- 0.002 | 0.961 +/- 0.003 | 0.0065 +/- 0.002 | 0.987 +/- 0.002 |
| MODL | 0.0071 +/- 0.004 | 0.917 +/- 0.002 | 0.0072 +/- 0.002 | 0.889 +/- 0.002 |
| Deep Decoder | 0.0032 +/- 0.005 | 0.901 +/- 0.008 | 0.0079 +/- 0.002 | 0.908 +/- 0.002 |
| DIP | 0.0028 +/- 0.005 | 0.891 +/- 0.007 | 0.0084 +/- 0.002 | 0.912 +/- 0.002 |
| Our (MRI) | 0.0021 +/- 0.002 | 0.931 +/- 0.002 | 0.0065 +/- 0.002 | 0.919 +/- 0.002 |
| Our (MRI+ Nat) | 0.0017 +/- 0.001 | 0.938 +/- 0.002 | 0.0060 +/- 0.002 | 0.923 +/- 0.002 |
| Our (MRI+Nat+Fractal) | 0.0012 +/- 0.002 | 0.940 +/- 0.002 | 0.0058 +/- 0.002 | 0.926 +/- 0.002 |

**TABLE 2.** fastMRI Brain: Our method outperformed all SOTA methods when trained on procedural images, i.e. fractral images, natural images from ImageNet, and a small set, i.e. only 20 samples from MRI database.

|  | R = 4 |  | R = 8 |  |
| --- | --- | --- | --- | --- |
|  | RMSE | SSIM | RMSE | SSIM |
| GRAPPA | 0.0199 +/- 0.002 | 0.799 +/- 0.002 | 0.0212 +/- 0.002 | 0.612 +/- 0.002 |
| SENSE | 0.0110 +/- 0.002 | 0.801 +/- 0.001 | 0.0235 +/- 0.002 | 0.620 +/- 0.002 |
| RAKI | 0.0138 +/- 0.006 | 0.901 +/- 0.002 | 0.0201 +/- 0.002 | 0.881 +/- 0.004 |
| JMODL | 0.0026 +/- 0.002 | 0.967 +/- 0.003 | 0.061 +/- 0.002 | 0.887 +/- 0.002 |
| MODL | 0.0053 +/- 0.004 | 0.952 +/- 0.002 | 0.0078 +/- 0.002 | 0.804 +/- 0.002 |
| Deep Decoder | 0.0072 +/- 0.005 | 0.949 +/- 0.008 | 0.0083 +/- 0.002 | 0.828 +/- 0.002 |
| DIP | 0.0061 +/- 0.005 | 0.971 +/- 0.007 | 0.0082 +/- 0.002 | 0.821 +/- 0.002 |
| Our (MRI) | 0.0037 +/- 0.002 | 0.974 +/- 0.002 | 0.0045 +/- 0.002 | 0.849 +/- 0.002 |
| Our (MRI + Nat) | 0.0031 +/- 0.001 | 0.978 +/- 0.002 | 0.0040 +/- 0.002 | 0.893 +/- 0.002 |
| Our (MRI + Nat + Fractal) | 0.0029 +/- 0.002 | 0.980 +/- 0.002 | 0.0038 +/- 0.002 | 0.906 +/- 0.002 |

### 3.3 Quantitative Results

Quantitative results are shown in Tables 2 and 1 for RMSE and SSIM scores. It can observed from Figs 5 and 4 difference images of GRAPPA, SENSE, RAKI, JMODL, VarNet, ConvDecoder, and Deep Image Prior (DIP) shows that our method is on par and most of the times our method is better than the SOTA methods. The SSIM and RMSE scores given below are averaged over 200 different mid-slice images from the FastMRI validation set.

#### Ablation Studies

We studied three types of inputs to the Eqn. 7 to learn the supermasks, i.e. (i) learn the supermask using the kspace measurements of MRI images; (ii) use kspace measurements of natural images of ImageNet dataset and only 20 samples of MRI kspace measurements to finetune; and the (iii) use a combination of fractal image kspace measurements and natural image kspace measurements from ImageNet, i.e. 50k fractal R = 2 undersampled kspace and 20k natural R = 2 undersampled kspace measurements, and learn the supemask using Eqn. 7. We then use only 20 MRI kspace measurements to finetune. The results are show in Table 2 and 1. Please note that of all cases we use the self supervised learning approach that we discussed in Sec. 2.2 and in Eqn. 7. We note that the choice of combination of fractal image kspace measurements and natural image kspace measurements from ImageNet outperforms all the SOTA methods.

## 4 DISCUSSION AND CONCLUSION

This work demonstrates a novel method to solve the MR reconstruction task without ever updating the weights of the network. We ask ourselves the question: “can a random weighted deep network structure encode informative cues to solve the MR reconstruction problem from highly under-sampled k-space measurements?” Our proposed methodology selects an optimal subnetwork from a randomly weighted dense network to perform MR reconstruction without updating the weights - neither at training time nor at inference time. The methodology does not require ground truth data and shows excellent performance across domains in T1-weighted (head, knee) images from highly under-sampled multi-coil k-space measurements.

Trained deep networks such as Variational Network and J-Model outperform classical MR reconstruction approaches (e.g. GRAPPA, SENSE) for highly under-sampled k-space data. However, they require large datasets at training time (along with ground truth data) which may not be readily available in practice. It is also observed that such methods are not task-agnostic and perform poorly for out-of-distribution test cases such as the reconstruction of brain images from a network trained on the knee dataset. Untrained networks such as DIP, DeepDecoder, ConvDecoder on the other hand do not require explicit training but perform optimization during inference time which makes them extremely slow, requiring several minutes to process a single slice of k-space data.

In this work, we address both these challenges and propose to perform fast multi-coil MRI reconstruction using a weight agnostic randomly weighted network (WAN) without the need for ground truth data. By design, the proposed algorithm is agnostic to different tasks in terms of SSIM, PSNR, and RMSE scores and generalizes better to out-of-distribution examples. The proposed method provides performance similar to highly trained networks, but without requiring ground truth data or abundant training datasets. The method is also very fast and can reconstruct data in real-time. The proposed methodology offers a novel perspective to learn the mapping from under-sampled k-space to the MR image. WAN is based on the premise that a subnetwork of a very large deep neural network can encode meaningful information to solve a task even though the parameters of the network are totally random. The proposed methodology does not require any ground truth MR image (or k-space measurements) and can perform reconstruction tasks in a domain agnostic manner. We show that the proposed method outperforms the state-of-the-art (SOTA) methods such as ConvDecoder, DIP, VarNet, etc., on the fastMRI knee and brain dataset.

Our future work includes pondering to the understand several aspects of the proposed methodology such as (i) correct reconstruction of minute details of pathology and anatomical structures; (ii) risk quantification; (iii) robustness; (iv) running time complexity; and (v) generalization.

## Abbreviations

MRI: Magnetic Resonance Imaging
WAN: weight agnostic randomlyweighted network
SGD: stochastic gradient descent

## Footnotes

1 Below are the official implementations of various methods we discussed: **VarNet**: https://github.com/VLOGroup/mri-variationalnetwork/ **JMODL**: https://github.com/hkaggarwal/J-MoDL.git **RAKI**: https://github.com/geopi1/DeepMRI.git **SENSE**: https://mrirecon.github.io/bart/ **GRAPPA**: https://github.com/mckib2/pygrappa.git **ConvDecoder**: https://github.com/MLI-lab/ConvDecoder.git

